# Human buccal epithelial microplicae maintain a characteristic wavelength despite variable network topology

**DOI:** 10.64898/2026.08.06.743190

**Authors:** Gail McConnell

**Author notes:** **Statements and Declarations:** The author has no conflicts of interest.

## Abstract

Microplicae are ridge-like membrane projections that are prominent features of many epithelial surfaces, yet little is known about the principles governing their spatial organisation. Determining whether microplicae represent stochastic membrane folds or biologically organised surface architectures is essential for understanding their formation and functional roles. Here, differential interference contrast images of human buccal epithelial cells were analysed using quantitative image-processing approaches. Ridge networks were segmented and characterised using complementary measurements of characteristic wavelength, including medial-axis and nearest-neighbour Voronoi analyses, together with skeleton-based metrics describing network architecture. Analysis of n=100 buccal epithelial cells sampled from n=10 donors revealed a reproducible sub-micron characteristic wavelength. Mean medial-axis spacing was 0.511 ± 0.042 µm and mean Voronoi nearest-neighbour spacing was 0.588 ± 0.057 µm. Characteristic wavelength exhibited CV of between only 8.26% and 9.67% across the dataset. However, metrics describing network architecture, including ridge density, branching and connectivity, varied by up to 109%. Donor-level analysis reported the same overall trends, with conservation of the characteristic wavelength while network parameters had considerably greater variation. These findings identify a previously unrecognised organising principle of microplical architecture, suggesting that epithelial membrane organisation is regulated through conservation of an intrinsic geometric length scale while network topology remains comparatively free to remodel.

## 1. Introduction

Microplicae are ridge-like projections of the apical plasma membrane that are a characteristic feature of many epithelial cell surfaces (Andrews, 1976). They are prominent on stratified squamous epithelia, including the corneal, conjunctival and oral mucosa, where they contribute substantially to the three-dimensional structure of the cell surface (Asikainen et al., 2015) (Landay and Schroeder, 1979). Although microplicae have been recognised through electron microscopy and more recently by high-resolution light microscopy (Lam et al., 2015), their biological function and the mechanisms governing their organisation remain poorly understood (Depasquale, 2018; Lu et al., 2022). Proposed roles include increasing membrane surface area, enhancing adhesion of mucus or tear film components, facilitating mechanical resilience, and organising membrane-associated proteins (Pinto et al., 2019). These proposed functions depend not only on the presence of microplicae but potentially also on how they are spatially organised across the epithelial surface. Consequently, determining whether microplical architecture reflects a biologically regulated structural organisation or simply the accumulation of irregular membrane folds represents an important unresolved question in epithelial cell biology. However, these hypotheses have largely been based on qualitative observations of surface morphology rather than quantitative analyses of architecture (Asikainen et al., 2015, 2014).

The morphology of microplicae has traditionally been described as an irregular, labyrinthine network of membrane ridges (Andrews, 1976; Asikainen et al., 2015). However, many apparently irregular biological patterns nevertheless exhibit highly conserved statistical organisation. Examples include vascular networks (Contessa and Luca, 2013), collagen matrices (Fratzl, 2008), trabecular bone (Callens et al., 2021), neuronal dendritic branching (Cuntz et al., 2010) and epithelial cell packing (Gibson et al., 2006), all of which display considerable local variability while maintaining reproducible global geometric properties. Such systems are increasingly understood through quantitative descriptors that characterise distributions, spatial frequencies and topological organisation rather than relying solely on qualitative morphology. Applying similar approaches to microplicae raises the possibility that the visually complex ridge networks observed on epithelial surfaces may likewise possess conserved structural features that are not immediately apparent by inspection.

One candidate organising principle is characteristic wavelength. In many physical and biological systems, labyrinthine patterns emerge through self-organising processes that generate structures with a preferred spatial frequency. Examples include reaction-diffusion systems (Kondo and Miura, 2010; Turing, 1952), elastic instabilities (Cerda and Mahadevan, 2003), phase separation (Bray, 1994), and active cytoskeletal processes (Lam et al., 2015; Lu et al., 2022; Pinto et al., 2019; Salbreux et al., 2012), all of which produce networks whose local geometry varies substantially while maintaining a relatively constant spacing between neighbouring ridges. If microplicae arise through analogous mechanisms, ridge spacing may represent a biologically regulated architectural parameter that reflects the physical processes governing epithelial surface organisation, whereas network topology may remain comparatively free to remodel. Conversely, if microplicae arise predominantly through stochastic membrane deformations, no conserved spatial scale would be expected either within or between cells. Demonstrating the existence of a conserved characteristic wavelength would therefore provide evidence that epithelial surface organisation is constrained by an underlying geometric organising principle rather than representing an arbitrary arrangement of membrane folds.

Despite the importance of this question, quantitative analyses of microplical organisation remain relatively limited (Asikainen et al., 2015; Depasquale, 2018). Recent work has demonstrated how advanced computational approaches can provide important insights into the dynamics and mechanics of microridge pattern formation (Bhavna and Sonawane, 2023). However, these approaches have not addressed whether microplical organisation is governed by a conserved geometric length scale. Previous quantitative studies have instead focused on descriptors such as ridge length, ridge density, branching and area coverage (Asikainen et al., 2015; Bale and White, 1982; Grossman, 1987). Although these metrics quantify aspects of network morphology and complexity, they do not directly measure the spacing between neighbouring ridges and therefore cannot determine whether a characteristic wavelength exists. Distinguishing geometric spacing from network topology therefore requires analytical approaches specifically designed to quantify these complementary aspects of architecture independently.

Human buccal epithelial cells provide an accessible model system for investigating microplical organisation (Groeger and Meyle, 2019). These cells possess prominent microplicae (Asikainen et al., 2015) and can be obtained non-invasively from multiple donors to test whether conserved architectural principles persist across individual cells and donors. Differential interference contrast microscopy offers excellent contrast (Nomarski, 1955; Smith, 1955) for visualising surface ridges without the need for fluorescent labelling, making it well suited to large-scale quantitative analyses of living epithelial cells.

In the present study, the hypothesis was tested that microplicae are organised around a conserved characteristic wavelength rather than representing an irregular collection of membrane folds. Differential interference contrast (DIC) images of live human buccal epithelial cells were analysed using an automated classical image-processing pipeline that combined ridge enhancement, segmentation and skeletonisation with complementary measurements of ridge spacing based on medial-axis and Voronoi nearest-neighbour analyses. These geometric measurements were integrated with quantitative descriptors of network topology, including ridge density, branching, connectivity and loop formation, to determine whether spatial wavelength and architectural complexity are independently regulated. Analysis reveals that, despite substantial variability in network architecture between cells, microplicae exhibit a remarkably conserved sub-micron characteristic wavelength. These findings identify a previously unrecognised organising principle of epithelial surface architecture and suggest that regulation of spatial wavelength may represent a fundamental constraint governing microplical organisation.

## 2. Methods

### 2.1. Specimen preparation

Human buccal epithelial cells were collected from n=10 healthy adult donors using sterile cotton-tipped swabs. Cells were transferred immediately to a clean glass microscope slide and gently dispersed to produce a sparse monolayer of isolated cells. A Type 1.5 glass coverslip was carefully placed over the specimen to form a thin layer suitable for high-resolution transmitted-light imaging. Samples were imaged immediately following preparation to preserve native cell morphology.

### 2.2. Differential interference contrast microscopy of cell specimens

DIC microscope imaging was performed with an upright light microscope (Eclipse Ni, Nikon) equipped with a 100x/1.45 numerical aperture oil immersion lens (Plan Apo λ, Nikon). Images with an 8-bit depth were acquired with a monochromatic camera (CellCam Centro 200MR, Cairn Research) controlled by MicroManager (v2.0.0) (Edelstein et al., 2014). For each specimen provided by the n=10 donors, n=10 images were acquired, with a minimum of one cell per image. All data were saved in the OME.TIFF format.

### 2.3. Image processing and image data analysis

Microplicae organisation was quantified using a custom Python image-analysis pipeline designed to test whether epithelial surface ridges exhibit a conserved characteristic wavelength. Differential interference contrast images were processed using an automated analysis pipeline comprising image normalisation, ridge enhancement, segmentation, skeletonization, and quantitative measurement of ridge spacing, and network topology. The major stages of the workflow are illustrated in Figure 1.

**Fig. 1.**
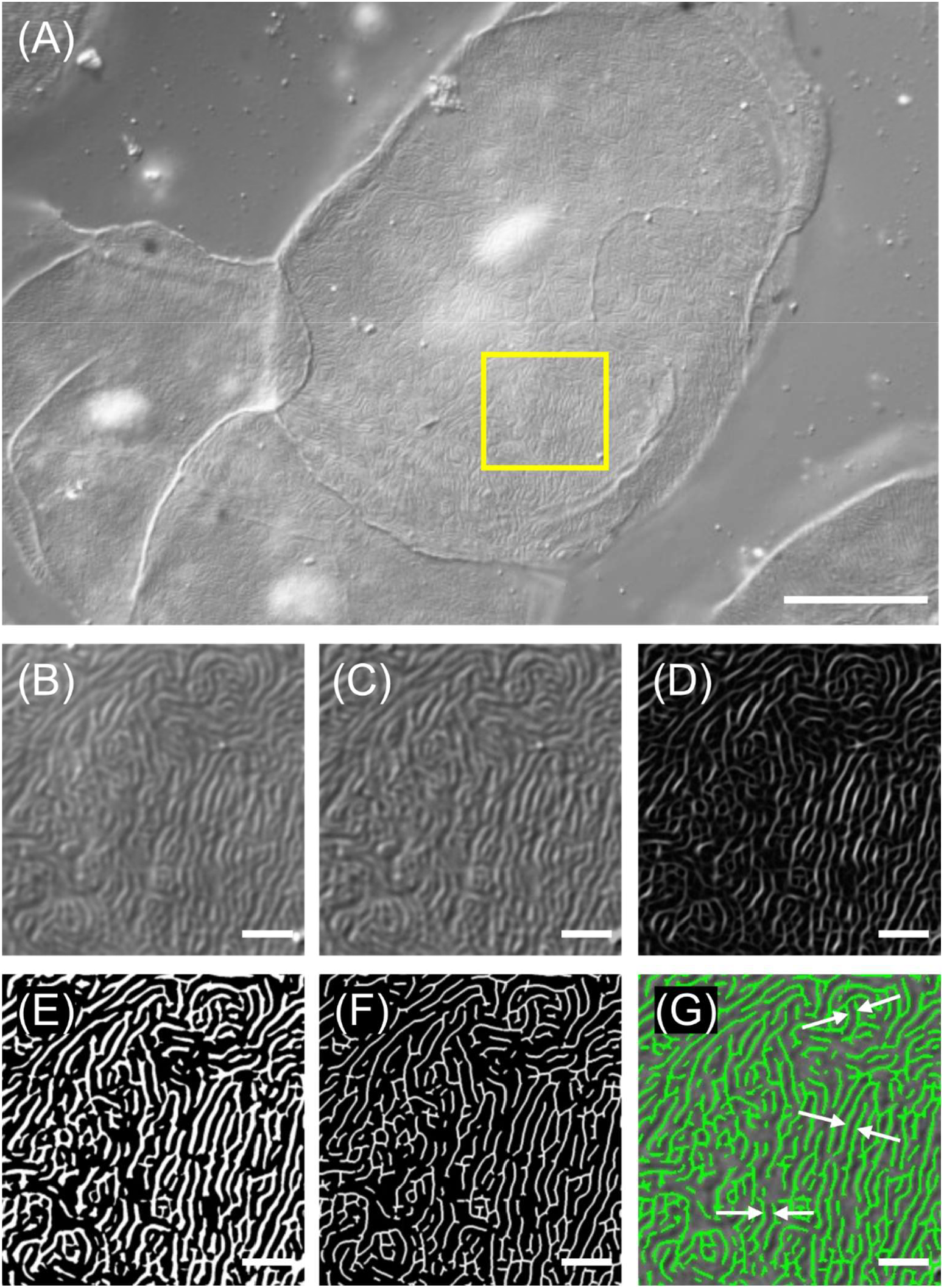
Image-processing pipeline used to quantify microplical architecture from differential interference contrast (DIC) images of human buccal epithelial cells. (A) Representative DIC image showing the labyrinthine microplical network on the apical surface of a buccal epithelial cell. The yellow box indicates the region of interest shown in panels B-G. (B) High-pass pre-processed image after intensity normalisation and background subtraction. (C) Ridge-enhanced image following application of the Sato ridge filter. (D) Binary ridge segmentation obtained by Otsu thresholding. (E) Skeletonised representation of the segmented ridge network used for quantitative analysis of topology and spacing. (F) Cleaned skeleton following removal of short, disconnected ridge components. (G) Skeleton overlay in green. Example ridge spacings (white arrows) identified for network topology analysis. Scale bars: (A) 10 μm; (B-G) 1 μm.

The image-analysis pipeline was implemented in Python and executed on a laptop computer running Microsoft Windows 11 Enterprise (version 10.0.26200 Build 26200) on an x64-based architecture. The system was equipped with an Intel® Core™ Ultra 5 135U processor (12 cores, 14 logical processors) and 16 GB RAM. GPU acceleration was not used during image processing or analysis. The software was developed using open-source Python libraries including NumPy (Harris et al., 2020), SciPy (Virtanen et al., 2020), scikit-image (van der Walt et al., n.d.), pandas, Matplotlib, and Pillow for image processing, quantitative analysis, visualisation and data export.

The analysis script was written to process batches of microscopy images and to quantify ridge spacing and ridge-network architecture from each cell image. Images were imported as greyscale arrays and normalised to the full intensity range before analysis. Intensity values were converted to floating-point format, background-subtracted, and rescaled prior to ridge detection.

Differential interference contrast images were pre-processed to enhance fine surface ridges and suppress low-frequency illumination gradients. Pixel intensities were clipped between the 1^st^ and 99^th^ percentiles, rescaled, and then subjected to Gaussian background subtraction using σ=30 pixels. The resulting high-pass image was standardised and rescaled to a 0-1 intensity range, producing a binary image output.

Microplicae were detected as bright ridge-like structures using a Sato ridge filter (Sato et al., 1998). The ridge-enhanced image was thresholded using Otsu’s method (N Otsu, 1979), and binary ridge masks were then skeletonised to generate one-pixel-wide representations of the microplical network for spacing and topology analysis.

Characteristic ridge spacing was measured using two complementary approaches. First, a medial-axis method was applied to the inverse skeleton image. The Euclidean distance transform was calculated for the inter-ridge space, and the medial axis of this background region was used to identify positions equidistant from adjacent ridge skeletons (Blum, 1967). Local ridge-to-ridge spacing was calculated as twice the distance from each medial-axis point to the nearest skeleton pixel. Measurements within 0.5 μm of the image border were excluded, and spacing values were retained only if they fell between 0.20 μm and 1.20 μm.

Ridge spacing was also independently measured using a Voronoi nearest-neighbour approach (Aurenhammer, 1991). Skeleton components shorter than 0.15 μm (which corresponds to sub-diffraction-limited features) were removed, and the remaining skeleton was labelled into connected ridge components. Adjacent Voronoi regions were identified from neighbouring label boundaries, and for each adjacent ridge pair the minimum nearest-neighbour distance between the two skeleton components was calculated using a k-d tree search. Pairwise distances were retained only if they were between 0.20 μm and 1.20 μm.

For each image, global spacing distributions were summarised by the number of measurements, mean, median, standard deviation, coefficient of variation (CV), and 5^th^, 25^th^, 75^th^, and 95^th^ percentiles. The primary wavelength outputs used for downstream analysis were mean medial-axis spacing and mean Voronoi nearest-neighbour spacing.

Network architecture was quantified from the binary ridge mask and skeleton. Ridge area fraction was calculated as the fraction of image pixels occupied by the binary ridge mask. Skeleton length density was calculated as total skeleton length divided by image area. Endpoints were identified as skeleton pixels with one neighbouring skeleton pixel, whereas branch points were identified as skeleton pixels with three or more neighbours. Endpoint object density and branch-point object density were expressed per μm^2^. Network loop density was estimated from the Euler number and the number of connected ridge components, then normalised to image area.

The pipeline saved per-cell diagnostic images including the original greyscale image, pre-processed high-pass image, Sato ridge-enhanced image, binary ridge mask, skeleton, and skeleton overlay. Per-cell CSV files were also exported for detailed medial-axis spacing measurements, Voronoi nearest-neighbour ridge pairs, and network-level metrics.

### 2.4 Statistical analysis

Statistical analyses were performed using GraphPad Prism (v8.0.2). For each image, the automated image-analysis pipeline measured mean medial-axis spacing, mean Voronoi nearest-neighbour spacing, ridge area fraction, skeleton length density, endpoint object density, branch-point object density, and loop density.

The primary analysis was performed at the cell level, using all images (n = 100). Descriptive statistics are presented as the mean, standard deviation (SD), and 95% confidence interval (95% CI) unless otherwise stated. Variability in characteristic wavelength and network architecture across the complete dataset was quantified using the CV: this was calculated from the SD divided by the mean and was expressed as a percentage. These cell-level analyses were used to quantify the overall distribution and variability of microplical organisation within the study population.

The cell constituted the unit of observation for quantifying within-population variability in microplical organisation. However, because multiple cells were sampled from each donor, donors represented the independent biological replicates. Consequently, cell-level analyses were used to describe the distribution and variability of microplical architecture across individual cells, whereas donor-level analyses (n = 10 donors) were used to assess biological reproducibility between individuals. Inferential conclusions regarding biological reproducibility are therefore based primarily on the donor-level analysis.

To assess the reproducibility of these measurements between individuals, a donor-level analysis was also performed. For each donor (n = 10), the mean value of each quantitative parameter was calculated from ten images per donor, producing one representative value per donor. Donor-level descriptive statistics were calculated from these donor means to determine the extent of inter-individual variation in characteristic wavelength and network architecture.

All statistical analyses and decision thresholds were specified before analysis. Descriptive statistics constituted the primary outcome measures. Correlation analyses were exploratory and were evaluated using Pearson’s or Spearman’s correlation, as appropriate following assessment of normality by the Shapiro-Wilk test (Shapiro and Wilk, 1965). Statistical significance was assessed using a two-sided significance level of α = 0.05.

## 3. Results

### 3.1 Microplicae in human buccal epithelial cells have a highly conserved characteristic wavelength

Analysis of n=100 images of buccal epithelial cells obtained from n=10 healthy donors demonstrated that microplicae possess a reproducible sub-micron characteristic wavelength. Figure 2A compares the distributions obtained using the two independent spacing measurements, whereas Figure 2B demonstrates that characteristic wavelength is markedly less variable than network topology across the dataset.

**Fig. 2.**
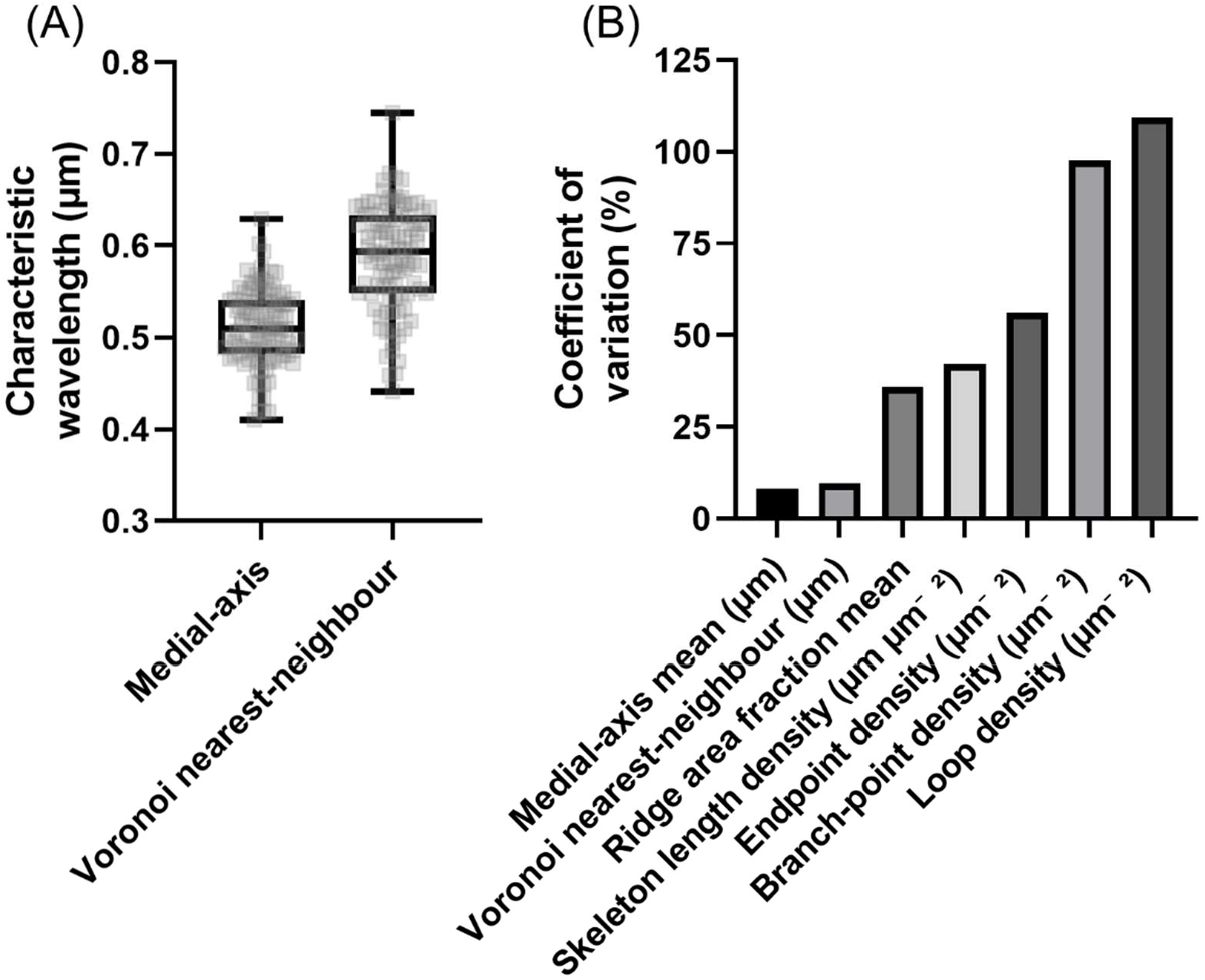
Comparison of variability in characteristic wavelength and network topology across all analysed cell images (n=100). (A) Distribution of characteristic wavelength measured using two independent approaches, namely medial-axis ridge spacing and Voronoi nearest-neighbour ridge spacing. The distribution of per-cell values is shown, with box plots indicating the median and interquartile range. (B) Coefficient of variation (CV) for characteristic wavelength measurements and network topology descriptors calculated across the complete dataset. Characteristic wavelength exhibited markedly lower variability, with CV∼8-10%, while measurements describing network architecture, including ridge area fraction, skeleton length density, endpoint density, branch-point density and loop density showed greater variation.

The mean ridge spacing given by the medial-axis analysis was 0.511 μm (SD 0.042 μm; 95% CI 0.502-0.519 μm), whereas the independent Voronoi nearest-neighbour analysis yielded a mean spacing of 0.588 μm (SD 0.057 μm; 95% CI 0.577-0.599 μm).

Although the two analytical methods differ in their geometric definitions of inter-ridge spacing, both produced consistent estimates of the underlying spatial scale of the microplical network. The systematic difference between the absolute values obtained by the two methods was expected because the medial-axis analysis measures local separation throughout the inter-ridge space, whereas the Voronoi analysis measures the minimum distance between neighbouring ridge skeletons.

Characteristic wavelength exhibited relatively little variability across the pooled dataset. The CV was 8.26% for medial-axis measurements and 9.67% for Voronoi nearest-neighbour measurements, indicating that ridge spacing is largely conserved.

### 3.2. Network topology varies substantially across different human buccal epithelial cells

In contrast to the conserved characteristic wavelength of microplicae, measurements describing ridge-network architecture exhibited greater variability across the dataset. Ridge area fraction had a mean value of 0.087 (SD 0.031; 95% CI 0.081-0.093), indicating that approximately 8.70% of the cell surface was occupied by segmented ridge structures. Skeleton length density averaged 0.461 μm μm^−2^ (SD 0.194 μm μm^−2^; 95% CI 0.422-0.499 μm μm^−2^).

Measurements describing network connectivity were considerably more variable. Endpoint object density averaged 1.069 μm^−2^ (SD 0.601 μm^−2^; 95% CI 0.950-1.189 μm^−2^), while branch-point object density averaged 0.497 μm^−2^ (SD 0.486 μm^−2^; 95% CI 0.401-0.594 μm^−2^). Loop density averaged 0.014 μm^−2^ (SD 0.015 μm^−2^; 95% CI 0.011-0.017 μm^−2^).

The relative variability of these topological descriptors was markedly greater than that of characteristic wavelength. CVs were 36.04% for ridge area fraction, 42.18% for skeleton length density, 56.19% for endpoint density, 97.64% for branch-point density and 109.3% for loop density.

Collectively, these pooled data demonstrate that the measured network topology features are substantially more variable than measurements of characteristic wavelength, suggesting that ridge spacing is more tightly conserved than network topology.

### 3.3 Characteristic wavelength is highly reproducible across donors

To determine whether the conserved characteristic wavelength reflected reproducibility between individuals rather than pooling across all analysed cells, donor-level summary statistics were calculated from ten images acquired from each of the ten donors (Figure 3). Both the medial-axis and Voronoi analyses demonstrated highly consistent characteristic wavelengths between donors. Mean medial-axis spacing was 0.511 μm (SD 0.025 μm; 95% CI 0.493-0.529 μm), corresponding to a CV of 4.90%. Similarly, donor mean Voronoi nearest-neighbour spacing was 0.588 μm (SD 0.036 μm; 95% CI 0.562-0.614 μm), with a CV of 6.12%.

**Fig. 3.**
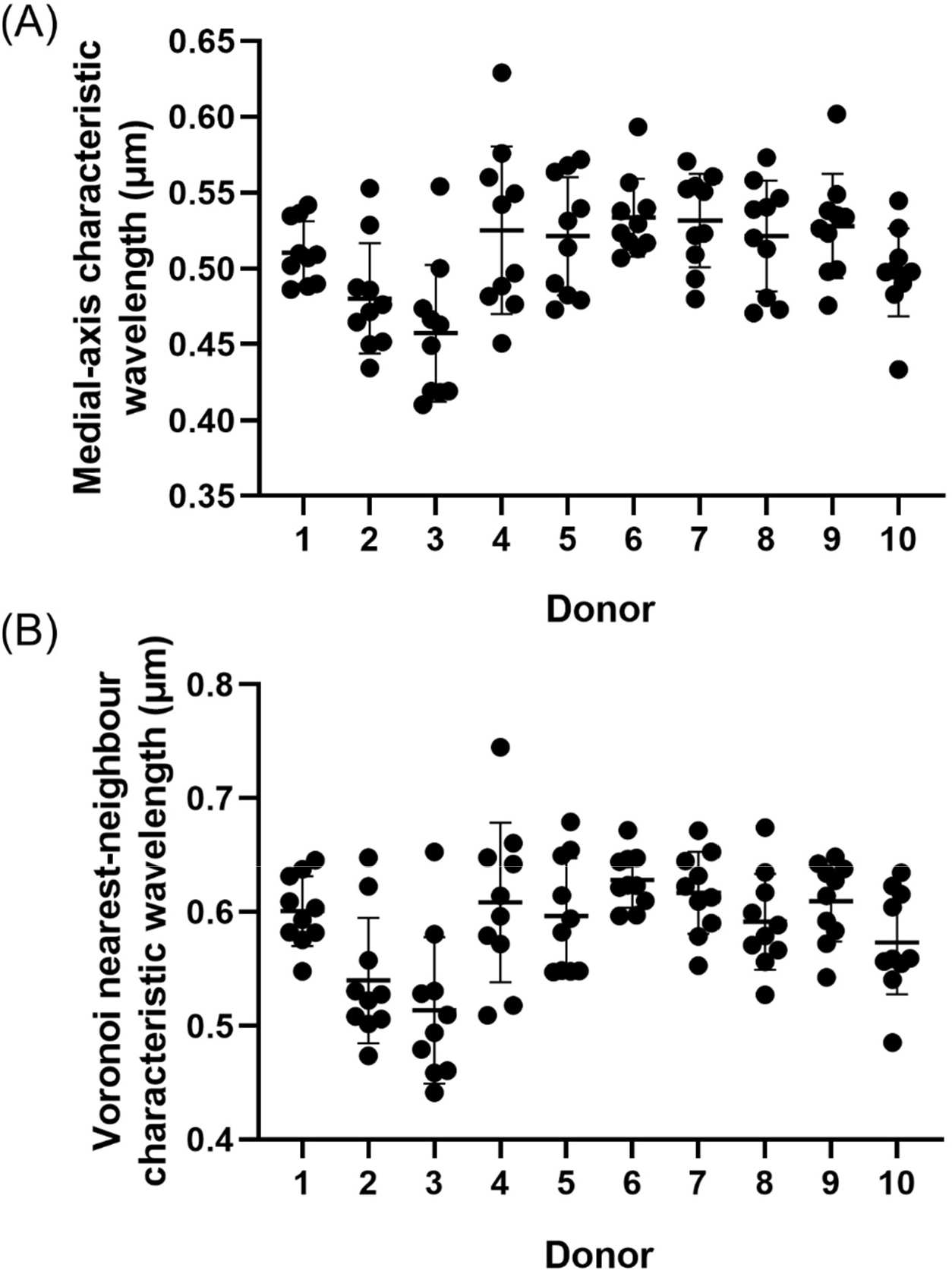
Donor-level reproducibility of characteristic wavelength measurements. (A) Mean medial-axis characteristic wavelength measured from ten cells for each donor (n=10 donors). Points represent individual cells, while horizontal bars indicate the donor mean ± SD. (B) Corresponding Voronoi nearest-neighbour characteristic wavelength measurements for the same donors. Although modest differences between donors were observed, the characteristic wavelength remained highly consistent across individuals, supporting the existence of a conserved geometric length scale in buccal epithelial microplicae.

Donor-level measurements of network architecture exhibited lower variability than the pooled cell-level analysis but remained substantially more variable than characteristic wavelength. Ridge area fraction averaged 0.087 (SD 0.017; 95% CI 0.075-0.099; CV 19.14%), skeleton length density averaged 0.461 μm μm^−2^ (SD 0.101 μm μm^−2^; 95% CI 0.388-0.533 μm μm^−2^; CV 21.94%), and endpoint object density averaged 1.069 μm^−2^ (SD 0.287 μm^−2^; 95% CI 0.865-1.274 μm^−2^; CV 26.79%). Branch-point object density averaged 0.497 μm^−2^ (SD 0.276 μm^−2^; 95% CI 0.300-0.695 μm^−2^; CV 55.50%), while loop density averaged 0.014 μm^−2^ (SD 0.008 μm^−2^; 95% CI 0.008-0.019 μm^−2^; CV 56.87%).

These donor-level analyses confirm that the conserved characteristic wavelength is reproducible across independent individuals. Although averaging measurements within donors reduced the variability of all quantitative descriptors, characteristic wavelength remained substantially more conserved than network topology. Even at the donor level, CV values for geometric spacing remained below 7%, whereas those for the most variable topological descriptors exceeded 55%, supporting the conclusion that microplical organisation is characterised by a conserved geometric length scale rather than a fixed network architecture.

## 4. Discussion

The principal finding of this study is that human buccal epithelial microplicae possess a highly conserved characteristic wavelength despite substantial variation in network architecture. Analysis of 100 cells sampled from 10 independent donors demonstrated that individual cells exhibit a highly conserved characteristic wavelength. Donor-level analyses further showed that this conserved wavelength is reproducible across independent biological replicates. In contrast, descriptors of network topology, including ridge density, branching, termination and loop formation, exhibited substantially greater variability. These findings indicate that the defining architectural feature of microplicae is not the precise geometry of individual ridges, but the maintenance of a conserved geometric length scale expressed as a characteristic spacing between neighbouring ridges.

Historically, microplicae have been described qualitatively as irregular, labyrinthine membrane folds (Andrews, 1976; Asikainen et al., 2015; Depasquale, 2018). This interpretation is consistent with observations that microridges undergo continual remodelling of ridge length and branching while maintaining an organised surface pattern (Lam et al., 2015). The present findings extend this interpretation by showing that the apparent irregularity of individual ridge trajectories masks a highly conserved underlying spatial organisation. This decoupling of geometry and topology demonstrates that conservation of spatial organisation does not require conservation of network architecture, indicating that epithelial cells regulate geometric length scale rather than individual ridge morphology.

The distinction between wavelength and topology is likely to be biologically important (Lu et al., 2022). Topological descriptors such as ridge density, branch frequency and loop formation describe how the network is connected, whereas characteristic wavelength defines the spatial scale over which the network is organised (Cuntz et al., 2010). These properties need not be regulated by the same mechanisms. The present findings suggest that epithelial cells may regulate membrane architecture primarily at the level of geometric length scale, while allowing network topology to remain comparatively flexible. Such an organisational strategy would permit continual remodelling of the microplical network in response to local mechanical or physiological influences while preserving the spatial framework within which epithelial surface functions are maintained. This behaviour resembles that of many self-organising biological systems, in which local rearrangements occur without altering the fundamental length scale of the pattern (Cross and Greenside, 2009; Kondo and Miura, 2010).

The observation that donor-level variability was lower than cell-level variability further supports this interpretation. As expected, averaging measurements across n=10 cell images per donor reduced the variability of all quantitative descriptors, but the characteristic wavelength remained the least variable parameter at both levels of analysis. The conservation of wavelength across independent individuals therefore indicates that the observed spatial scale is unlikely to be a consequence of stochastic variation or sampling effects. Instead, it appears to represent an intrinsic organisational property of human buccal epithelial microplicae.

The present study does not identify the mechanisms responsible for establishing this conserved spatial scale, but several classes of process could plausibly generate such behaviour. These include cortical actomyosin mechanics, in which interactions between actin organisation, contractility and membrane tension generate characteristic pattern scales (Chugh and Paluch, 2018), or membrane mechanical constraints that favour a preferred ridge separation by minimising bending or elastic energy (Rombouts et al., 2023). Although these mechanisms differ substantially, they share the common prediction that an intrinsic spatial scale can emerge while individual ridge trajectories remain free to remodel, consistent with the present observations. If wavelength is biologically regulated, conservation of this spatial scale may also have functional consequences by regulating the dimensions of inter-ridge compartments that organise the glycocalyx and stabilise the overlying mucus layer, or by influencing molecular transport within the immediate pericellular environment (Kesimer et al., 2013). Future studies should therefore investigate wavelength itself as a biologically regulated parameter, rather than treating microplicae solely as irregular membrane folds arising from cortical actin.

This work also demonstrates the value of quantitative image analysis for investigating epithelial surface architecture. Previous studies have relied predominantly on qualitative observations, simple morphological descriptors, or deep-learning frameworks requiring extensive annotated training datasets for segmentation (Bhavna and Sonawane, 2023). By combining complementary measurements of ridge spacing with quantitative descriptors of network topology, the present classical approach distinguishes spatial organisation from network connectivity, providing an objective framework for analysing epithelial surface architecture. The close agreement between two independent measures of ridge spacing further increases confidence that the observed characteristic wavelength reflects an underlying biological feature rather than an artefact of a particular analytical approach.

Several limitations of this methodology should be acknowledged. The study examined microplicae from a single epithelial tissue type and therefore cannot determine whether the conserved wavelength identified here represents a general feature of epithelial microplicae or is specific to buccal epithelial cells. Similarly, although DIC microscopy provides excellent contrast for visualising surface ridges in living cells, it does not directly report the underlying cytoskeletal organisation responsible for the formation of microplicae. The present study was designed to identify geometric principles of organisation rather than the molecular mechanisms that generate them. Establishing the relationship between conserved wavelength and the actin cytoskeleton will require future multi-modal or correlative microscopy studies, for example combining quantitative image analysis with high-resolution fluorescence imaging or electron microscopy.

An additional limitation is that the present analysis is descriptive rather than mechanistic. Although the conserved characteristic wavelength is reported here, the processes responsible for establishing and maintaining this spatial scale remain unknown. Several candidate mechanisms may be envisaged, including reaction-diffusion processes, cortical actomyosin mechanics, membrane tension, or local mechanical feedback between neighbouring ridges. Distinguishing between these possibilities will require experimental perturbation of the cytoskeleton and assessment of the resulting changes in both wavelength and network topology.

In conclusion, this study identifies a previously unrecognised organising principle of human buccal epithelial microplicae. Rather than representing an arbitrary collection of membrane folds, microplicae exhibit a conserved characteristic wavelength that is maintained despite substantial variation in network architecture and reproducible across independent donors. These findings suggest that epithelial surface organisation is characterised by an intrinsic geometric length scale, with ridge topology representing a more flexible architectural property. More broadly, these findings suggest that regulation of geometric length scale may represent a previously unrecognised principle governing epithelial surface organisation, providing a foundation for future studies investigating how membrane mechanics, cytoskeletal organisation, and disease influence epithelial architecture. Should conservation of characteristic wavelength prove to be shared across multiple epithelial tissues, it may represent a general organising principle of apical membrane architecture rather than a feature unique to buccal cells.

## Acknowledgements

This work was funded in part by the Medical Research Council (MR/K015583/1), the Biotechnology and Biological Sciences Research Council (BB/Z51486X/1 and BB/X005178/1), and the Leverhulme Trust. I would like to thank Dr W.B. Amos for the loan of the differential interference microscope and camera used in this study.

## References

Andrews, P.M., 1976. Microplicae: characteristic ridge-like folds of the plasmalemma. Journal of Cell Biology 68, 420–429. 10.1083/jcb.68.3.420

Asikainen, P., Mikkonen, J.J., Ruotsalainen, T.J., Koistinen, A.P., Kullaa, A.M., 2014. Microstructure of the Superficial Epithelial Cells of the Human Oral Mucosa. Ultrastructural Pathology 38, 6–12. 10.3109/01913123.2013.816401

Asikainen, P., Sirviö, E., Mikkonen, J.J.W., Singh, S.P., Schulten, E.A.J.M., ten Bruggenkate, C.M., Koistinen, A.P., Kullaa, A.M., 2015. Microplicae – Specialized Surface Structure of Epithelial Cells of Wet-Surfaced Oral Mucosa. Ultrastructural Pathology 39, 299–305. 10.3109/01913123.2015.1054015

Aurenhammer, F., 1991. Voronoi diagrams—a survey of a fundamental geometric data structure. ACM Comput. Surv. 23, 345–405. 10.1145/116873.116880

Bale, E., White, F.H., 1982. Quantitative light and electron microscopical studies of the epithelial-connective tissue junction in intraoral mucosae. Journal of Microscopy 128, 69–78. 10.1111/j.1365-2818.1982.tb00438.x

Bhavna, R., Sonawane, M., 2023. A deep learning framework for quantitative analysis of actin microridges. npj Systems Biology and Applications 9, 21. 10.1038/s41540-023-00276-7

Blum, H., 1967. A transformation for extracting new descriptors of shape, Models for the Perception of Speech and Visual Form. MIT Press.

Bray, A.J., 1994. Theory of phase-ordering kinetics. Advances in Physics 43, 357–459. 10.1080/00018739400101505

Callens, S.J.P., Tourolle né Betts, D.C., Müller, R., Zadpoor, A.A., 2021. The local and global geometry of trabecular bone. Acta Biomaterialia 130, 343–361. 10.1016/j.actbio.2021.06.013

Cerda, E., Mahadevan, L., 2003. Geometry and Physics of Wrinkling. Physical Review Letters 90, 074302. 10.1103/PhysRevLett.90.074302

Chugh, P., Paluch, E.K., 2018. The actin cortex at a glance. Journal of Cell Science 131, jcs186254. 10.1242/jcs.186254

Contessa, P., Luca, C.J.D., 2013. Neural control of muscle force: indications from a simulation model. Journal of Neurophysiology 109, 1548–1570. 10.1152/jn.00237.2012

Cross, M., Greenside, H., 2009. Pattern Formation and Dynamics in Nonequilibrium Systems. Cambridge University Press, Cambridge. 10.1017/CBO9780511627200

Cuntz, H., Forstner, F., Borst, A., Häusser, M., 2010. One Rule to Grow Them All: A General Theory of Neuronal Branching and Its Practical Application. PLOS Computational Biology 6, e1000877. 10.1371/journal.pcbi.1000877

Depasquale, J.A., 2018. Actin Microridges. The Anatomical Record 301, 2037–2050. 10.1002/ar.23965

Edelstein, A.D., Tschudia, M.A., Amodaj, N., Pinkard, H., Vale, R.D., Stuurman, N., 2014. Advanced methods of microscope control using μManager software. Journal of Biological Methods 1, e10. https://doi.org/doi:%2010.14440/jbm.2014.36

Fratzl, P., 2008. Collagen: Structure and Mechanics. Springer.

Gibson, M.C., Patel, A.B., Nagpal, R., Perrimon, N., 2006. The emergence of geometric order in proliferating metazoan epithelia. Nature 442, 1038–1041. 10.1038/nature05014

Groeger, S., Meyle, J., 2019. Oral Mucosal Epithelial Cells. Frontiers in Immunology Volume 10-2019.

Grossman, E., 1987. A histometric/scanning electron microscope study of normal and loaded oral epithelium of the vervet monkey. J Anat 154, 81–90.

Harris, C.R., Millman, K.J., van der Walt, S.J., Gommers, R., Virtanen, P., Cournapeau, D., Wieser, E., Taylor, J., Berg, S., Smith, N.J., Kern, R., Picus, M., Hoyer, S., van Kerkwijk, M.H., Brett, M., Haldane, A., del Río, J.F., Wiebe, M., Peterson, P., Gérard-Marchant, P., Sheppard, K., Reddy, T., Weckesser, W., Abbasi, H., Gohlke, C., Oliphant, T.E., 2020. Array programming with NumPy. Nature 585, 357–362. 10.1038/s41586-020-2649-2

Kesimer, M., Ehre, C., Burns, K.A., Davis, C.W., Sheehan, J.K., Pickles, R.J., 2013. Molecular organization of the mucins and glycocalyx underlying mucus transport over mucosal surfaces of the airways. Mucosal Immunology 6, 379–392. 10.1038/mi.2012.81

Kondo, S., Miura, T., 2010. Reaction-Diffusion Model as a Framework for Understanding Biological Pattern Formation. Science 329, 1616–1620. 10.1126/science.1179047

Lam, P., Mangos, S., Green, J.M., Reiser, J., Huttenlocher, A., 2015. In Vivo Imaging and Characterization of Actin Microridges. PLOS ONE 10, e0115639. 10.1371/journal.pone.0115639

Landay, M.A., Schroeder, H.E., 1979. Differentiation in normal human buccal mucosa epithelium. Journal of Anatomy 128, 31–51.

Lu, T.Q., van Loon, A.P., Sagasti, A., 2022. How to wrinkle a cell: Emerging mechanisms of microridge morphogenesis. Current Opinion in Cell Biology 76, 102088. 10.1016/j.ceb.2022.102088

N Otsu, 1979. A Threshold Selection Method from Gray-Level Histograms. IEEE Transactions on Systems, Man, and Cybernetics 9, 62–66. 10.1109/TSMC.1979.4310076

Nomarski, G., 1955. Microinterféromètre différentiel à ondes polarisées. J. Phys. Rad. 16, 9S–13S.

Pinto, C.S., Khandekar, A., Bhavna, R., Kiesel, P., Pigino, G., Sonawane, M., 2019. Microridges are apical epithelial projections formed of F-actin networks that organize the glycan layer. Scientific Reports 9, 12191. 10.1038/s41598-019-48400-0

Rombouts, J., Elliott, J., Erzberger, A., 2023. Forceful patterning: theoretical principles of mechanochemical pattern formation. EMBO Reports 24, EMBR202357739. 10.15252/embr.202357739

Salbreux, G., Charras, G., Paluch, E., 2012. Actin cortex mechanics and cellular morphogenesis. Trends in Cell Biology 22, 536–545. 10.1016/j.tcb.2012.07.001

Sato, Y., Nakajima, S., Shiraga, N., Atsumi, H., Yoshida, S., Koller, T., Gerig, G., Kikinis, R., 1998. Three-dimensional multi-scale line filter for segmentation and visualization of curvilinear structures in medical images. Medical Image Analysis 2, 143–168. 10.1016/S1361-8415(98)80009-1

Shapiro, S.S., Wilk, M.B., 1965. An Analysis of Variance Test for Normality (Complete Samples). Biometrika 52, 591–611. 10.2307/2333709

Smith, F., 1955. Microscopic interferometry. Research 8, 385–395.

Turing, A.M., 1952. The chemical basis of morphogenesis. Philosophical Transactions of the Royal Society of London. B, Biological Sciences 237, 37–72. 10.1098/rstb.1952.0012

van der Walt, S., Schoenberger, J., Nunez-Iglesias, J., Boulogne, F., Warner, J., Yager, N., Gouillart, E., Yu, T., n.d. scikit-image: image processing in Python. PeerJ 2, e453. 10.7717/peerj.453

Virtanen, P., Gommers, R., Oliphant, T.E., Haberland, M., Reddy, T., Cournapeau, D., Burovski, E., Peterson, P., Weckesser, W., Bright, J., van der Walt, S.J., Brett, M., Wilson, J., Millman, K.J., Mayorov, N., Nelson, A.R.J., Jones, E., Kern, R., Larson, E., Carey, C.J., Polat, İ., Feng, Y., Moore, E.W., VanderPlas, J., Laxalde, D., Perktold, J., Cimrman, R., Henriksen, I., Quintero, E.A., Harris, C.R., Archibald, A.M., Ribeiro, A.H., Pedregosa, F., van Mulbregt, P., Vijaykumar, A., Bardelli, A.P., Rothberg, A., Hilboll, A., Kloeckner, A., Scopatz, A., Lee, A., Rokem, A., Woods, C.N., Fulton, C., Masson, C., Häggström, C., Fitzgerald, C., Nicholson, D.A., Hagen, D.R., Pasechnik, D.V., Olivetti, E., Martin, E., Wieser, E., Silva, F., Lenders, F., Wilhelm, F., Young, G., Price, G.A., Ingold, G.-L., Allen, G.E., Lee, G.R., Audren, H., Probst, I., Dietrich, J.P., Silterra, J., Webber, J.T., Slavič, J., Nothman, J., Buchner, J., Kulick, J., Schönberger, J.L., de Miranda Cardoso, J.V., Reimer, J., Harrington, J., Rodríguez, J.L.C., Nunez-Iglesias, J., Kuczynski, J., Tritz, K., Thoma, M., Newville, M., Kümmerer, M., Bolingbroke, M., Tartre, M., Pak, M., Smith, N.J., Nowaczyk, N., Shebanov, N., Pavlyk, O., Brodtkorb, P.A., Lee, P., McGibbon, R.T., Feldbauer, R., Lewis, S., Tygier, S., Sievert, S., Vigna, S., Peterson, S., More, S., Pudlik, T., Oshima, T., Pingel, T.J., Robitaille, T.P., Spura, T., Jones, T.R., Cera, T., Leslie, T., Zito, T., Krauss, T., Upadhyay, U., Halchenko, Y.O., Vázquez-Baeza, Y., SciPy 1.0 Contributors, 2020. SciPy 1.0: fundamental algorithms for scientific computing in Python. Nature Methods 17, 261–272. 10.1038/s41592-019-0686-2

